# A genome-wide association study of high-sensitivity C-reactive protein in a large Korean population highlights its role in cholesterol metabolism

**DOI:** 10.1101/2024.06.27.600933

**Authors:** Kwangyeon Oh, Minju Yuk, Soyoun Yang, Jiyeong Youn, Qian Dong, Zhaoming Wang, Nan Song

**Affiliations:** Department of Pharmacy, College of Pharmacy, Chungbuk National University; Department of Epidemiology and Cancer Control, St. Jude Children’s Research Hospital, USA

**Author notes:** **Corresponding author** Nan Song, PhD. These authors contributed equally to this work (Minju Yuk, Soyoun Yang, Jiyeong Youn, Qian Dong). These authors also contributed equally to this work (Zhaoming Wang, Nan Song).

## Abstract

High-sensitivity C-reactive protein (hsCRP) is a representative biomarker of systemic inflammation and is associated with numerous complex diseases. To explore the biological pathways and functions underlying chronic inflammation, we conducted a genome-wide association study (GWAS) and several post-GWAS analyses of the hsCRP levels. This study was performed on data from 71,019 Koreans and is one of the largest East Asian studies. Overall, 69 independent single nucleotide polymorphisms (SNPs) were identified, including 12 novel variants located within *SHE, RP11-740C1.2, FCRL6, STEAP1B, AC002480.5, TOMM7, SPPL3, RP11-216P16.6, RP11-209K10.2, CTC-490E21.12, CYP2F2P, CBLC,* and *PVRL2*. The implicated genes and pathways are primarily involved in cholesterol metabolism and the immune response. A phenome-wide association study was performed based on a polygenic risk score constructed using 69 hsCRP-associated SNPs. Notably, the alleles associated with higher hsCRP levels appeared to be associated with lower low-density lipoprotein cholesterol levels (*P*=1.69 × 10^-33^, β=−1.47). Our findings provide evidence of a link between hsCRP and cholesterol as well as the clinical importance of hsCRP. Specifically, we suggest that genetically determined hsCRP levels may be useful for predicting the potential risk of cardiovascular or other diseases related to lipid metabolism.

**Author’s summary:** Chronic inflammation is associated with numerous complex diseases, including cancer, cardiovascular disease, and diabetes. Previous studies have shown that East Asians have lower levels of C-reactive protein (CRP), a biomarker of inflammation, than other ancestry groups. To identify East Asian-specific single nucleotide polymorphisms (SNPs) associated with chronic inflammation, we conducted a genome-wide association study and secondary genetic analyses on hsCRP in 71,019 Korean individuals. In total, 69 SNPs were identified, of which 12 were novel. A majority of the identified SNPs are located within genes (*LEPR*, *IL6R*, and *APOE*) that are involved in cholesterol metabolism and immune response. Notably, we found genetically determined hsCRP level may contribute to decrease cholesterol level, in contrast to previous epidemiological studies. Therefore, we suggest genetically determined hsCRP as a clinical tool for predicting the potential risk of abnormal cholesterol metabolism and its related diseases.

## Introduction

Chronic inflammation is associated with numerous complex diseases such as cancers, cardiovascular diseases (CVD), Alzheimer’s disease, autoimmune diseases, and metabolic syndrome (1–4). C-reactive protein (CRP), a key indicator and a clinical biomarker of systemic inflammation, is elevated in acute or chronic inflammatory conditions (5), and plays a biologically important role in mediating the inflammatory response (6). In conditions of inflammation, such as rheumatoid arthritis and infection, CRP is mainly produced by hepatocytes in the liver and released into the bloodstream under the control of IL-6 (6). Upon recognizing phosphocholine residues on pathogens, CRP activates the complement and opsonization systems, thereby promoting innate immunity (4). Additionally, CRP recognizes oxidized low-density lipoprotein (ox-LDL) along the blood vessels as a pathogen, triggering inflammation and innate immunity. CRP contributes to chronic inflammatory diseases (e.g., atherosclerosis, cancers, and type-2 diabetes) by increasing ox-LDL uptake by macrophages (4, 7).

Despite the clinical importance of CRP, it is challenging to conduct a genetic analysis of chronic inflammation using CRP levels because low CRP levels (<0.01 mg/L) are difficult to detect (8). However, with the development of a high-sensitivity CRP (hsCRP) assay, low CRP levels (<0.01 mg/L) can be detected, even in healthy individuals, thus predicting the risk of chronic inflammatory diseases (e.g., CVD, stroke, and myocardial infarction) (9). The heritability of hsCRP levels was high and estimated to be 44%–59%, indicating that genetic variation is a major determinant of hsCRP levels (10). Given the genetic importance of hsCRP, several genome-wide association studies (GWAS) have identified single nucleotide polymorphisms (SNPs) associated with hsCRP levels (rs1260326, 2p23.3, *GCKR*; rs147711004, 19q13.32, *BCAM - NECTIN2*; and rs4656241, 1q23.2, *OR10J6P - CRPP1*) (11). In addition, a GWAS identified hsCRP levels as a diagnostic biomarker for a monogenic diabetes, *HNF1A-*maturity-onset diabetes of the young (12, 13). However, previous GWAS studies have been predominantly conducted in individuals of European ancestry (11, 14). Differences in allelic architecture and linkage disequilibrium (LD) across ancestral populations could contribute to the heterogeneity of genetic susceptibility to hsCRP and related traits (15). This might partially explain why CRP levels vary among different ancestral populations; for example, serum CRP levels were reported to be lower in East Asians (EAs) than in Europeans (16). Although a previous GWAS was published with three SNPs (rs7553007, 1q23.2, CRP; rs2393791, 12q24.31, HNF1A; and rs9375813, 6q23.2, ARG1) associated with CRP levels in 8,529 Koreans (17), a new expanded GWAS of hsCRP levels with a much larger sample size may lead to the discovery of novel genetic susceptibility loci in an understudied EA population and the improved understanding of biological mechanisms and the clinical implication across populations.

GWAS or gene-/pathway-based analyses are likely to provide insights into the biological mechansims of diseases and traits (18). Functional genes involved in innate immunity (e.g., *CRP, IL6, IL6R, NLRP3,* and *IL1RN*) or metabolites (e.g., *APOC1, HNF1A, LEPR, GCKR, HNF4A, RORA,* and *PPP1R3B)* have been reported to be associated with hsCRP levels (11, 19). Furthermore, a phenome-wide association study (PheWAS) could provide a holistic view of associations between specific SNPs and a wide range of phenotypes, including diseases and lifestyle traits (20). Genetically determined hsCRP levels have been shown to be associated with dementia, hypercholesterolemia, and cancers (11, 21), suggesting either a potential causative link between hsCRP levels and these diseases or the existence of genetic pleiotropic effects.

Here, we leveraged the Korean Genome and Epidemiology Study (KoGES) with more than 70,000 participants to conduct one of the largest GWAS of hsCRP levels in an ancestry (EA) population. We further explored the potential underlying pathways involved in CRP regulation based on the identified hsCRP susceptibility SNPs, and further association between genetically determined hsCRP levels and some diseases and traits.

## RESULTS

### Characteristics of the study population

In this study, subjects were recruited from three population-based KoGES cohorts (22); Health examinee Study (HEXA), Cardiovascular Disease Association Study (CAVAS), and the Ansan/Ansung study. The epidemiology and clinical characteristics of the three cohort populations are summarized in **Table S1** (N_HEXA_=58,434 N_CAVAS_=7,108 N_Ansan/Ansung study_=5,477). Among the three KoGES cohorts, CAVAS subjects had the highest hsCRP levels (HEXA: 1.03 mg/L, CAVAS: 1.27 mg/L, Ansan/Ansung study: 1.08 mg/L, *P*<2×10^-16^) as well as the highest average age (HEXA: 54.7 years, CAVAS: 58.67 years, Ansan/Ansung study: 55.16 years). Among all the participants (N=71,019), 35.91% were men, 26.98% had history of smoking, 50.24% engaged in regular physical activity, and 22.11% were diagnosed with hypertension. The average (± SD) of age, body mass index (BMI), and hsCRP levels were 55.16 years (± 8.44), 24 Kg/m^2^ (± 2.93), and 1.06 mg/L (± 1.44) respectively.

Nineteen demographic variables were considered, 12 of which (sex, age, BMI, family income, alcohol drinking, regular physical activity, educational attainment, hypertension, diabetes, cardiovascular diseases, arthritis, and asthma) were significantly associated with hsCRP levels in all three KoGES cohorts (*P*<0.05) **(Table S2)**. As a result of backward selection of the 12 significant independent variables, six (sex, age, BMI, regular physical activity, smoking, and history of hypertension) were selected as covariates in the regression model for the GWAS analysis.

### GWAS on hsCRP levels

**Fig S3** presents the results of the GWAS on hsCRP for the three KoGES cohorts and **Fig S4** illustrates the results of the meta-analysis. In the current GWAS, we identified 69 independent hsCRP susceptibility SNPs (marked by the lead SNPs) near 41 genes (*P_meta_*<5×10^-8^), 12 of which were novel manifesting low LD with previously identified SNPs (r^2^<0.1) **(Fig 1, Data S1)**. The novel SNPs were rs117103761 (*SHE*), rs138233898 (*RP11-740C1.2*), rs189750399 (*FCRL6*), rs78821961 (*STEAP1B* and *AC002480.5*), rs74987345 (*Intergenic*), rs80348264 (*TOMM7*), rs149156910 (*SPPL3*), rs76847996 (*RP11-216P16.6*), rs1899754(*RP11-209K10.2*), rs28608119 (*RP11-209K10.2*), rs147400515 (*CTC-490E21.12* and *CYP2F2P*), rs145031198 (*CBLC*), and rs4599021 (*PVRL2*). The most significant three SNPs were rs3093068 mapped to *CRP* (β(C)_meta_=0.246, *P*_meta_=1.28×10^-304^), rs429358 mapped to *APOE* (β(C)_meta_=−0.224, *P*_meta_=7.21×10^-194^), and rs2393775 mapped to *HNF1A* (β(G)_meta_=−0.108, *P*_meta_=1.48×10^-121^). In this study, 10.15% of the total variance in hsCRP levels was explained by the lead SNPs.

**Fig 1.**
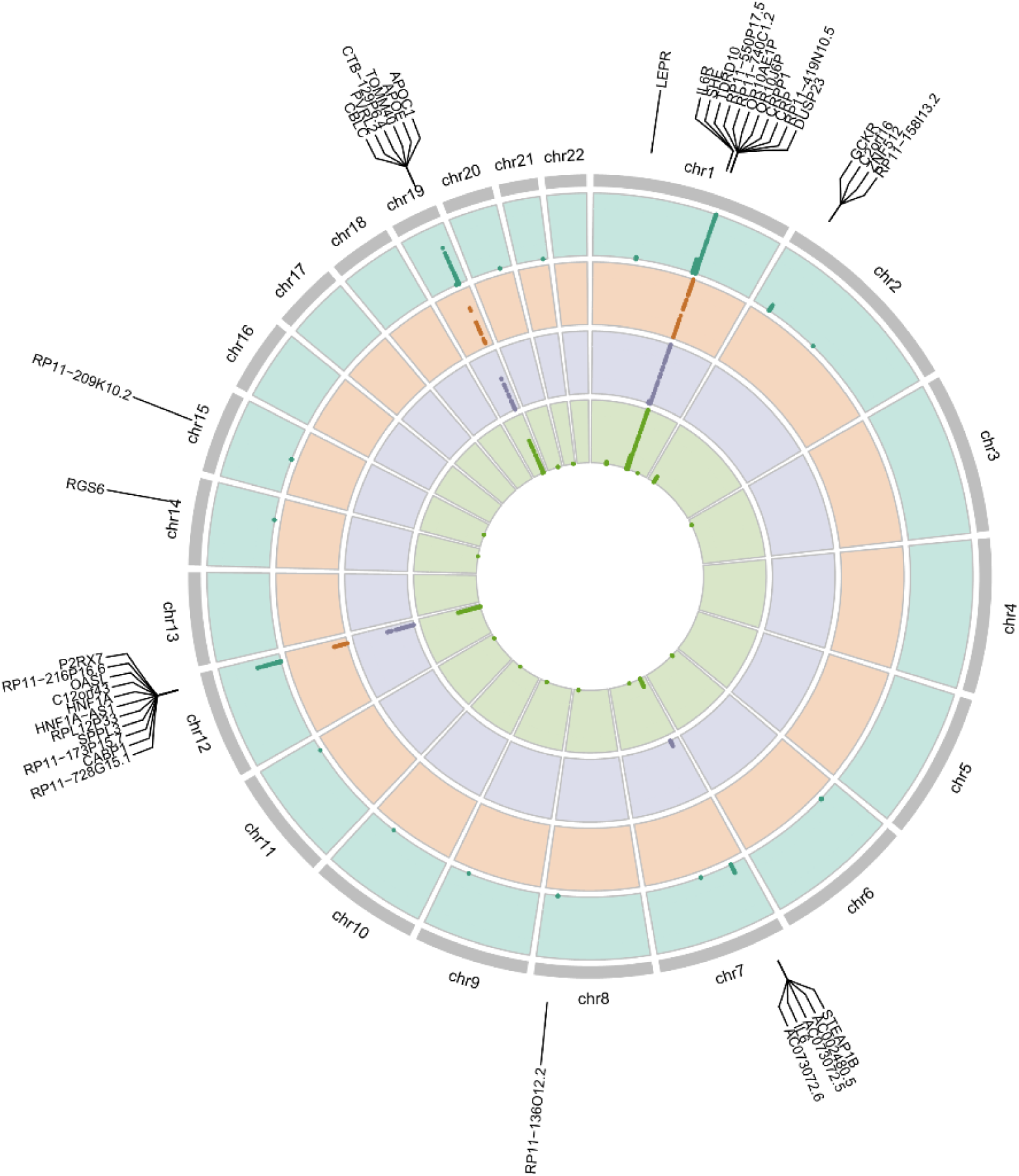
Circular Manhattan plot of GWAS on hsCRP. Circular Manhattan plot shows the genome-wide significant hits (*P*<5×10^-8^) of the GWAS results from the HEXA, CAVAS, and Ansan/Ansung study, and meta-analysis of the three cohort studies in order from the innermost part to the outermost part. Labelled genes were the nearest genes to the 85 lead SNPs.

Using our GWAS results, we evaluated the significance of hsCRP susceptibility SNPs identified by previously published GWAS. Validation analyses were conducted on 123 SNPs identified in EA ancestry populations and 260 and 239 SNPs identified in multi-ancestry populations with and without EA, respectively **(Fig S2)**. For the EA ancestry SNP group, 61 SNPs were validated, suggesting a validation rate of 51.5% (61/123) **(Data S2).** In contrast, the validation rates for multi-ancestry SNPs were relatively low, with 29.2% (76/260) of EA and 36.8% (88/239) without EA **(Data S3-4**).

### Functional analysis

Based on the hsCRP GWAS results, various functional analyses were conducted using the Functional Mapping and Annotation of GWAS (FUMA) (23), Multi-marker analysis of genomic annotation (MAGMA) (24), Search tool for the retrieval of interacting genes (STRING) (25), and ClueGO platforms (26). In our FUMA-ANNOVAR analysis, 46.5% of the significant hsCRP SNPs (*P*<5×10^-8^) were located in intronic regions, 35.4% in intergenic regions, and only 1.80% in exonic regions **(Fig S5)**. According to the MAGMA gene-based analysis, the SNPs were mapped to 18,681 protein-coding genes, of which 58 were significant with Bonferroni correction (*P*<2.68×10^-6^) **(Fig 2a, Table S3)**. The most significantly associated five genes were *LEPR* (*P*=1.38×10^-18^), *APOC1* (*P*=5.55×10^-17^), *OR10J1* (*P*=2.22×10^-16^), *MLEC* (*P*=2.28×10^-15^), and C2ofr16 (*P*=3.33×10^-15^). STRING protein-protein interaction analysis was conducted on 58 significant genes, resulting in the identification of complex interactions within 39 genes **(Fig 2b)**. In the MAGMA gene-property analysis, the abundance of hsCRP-associated genes expression was nominally significant in the liver (*P*=0.03) and blood (*P*=0.04) tissues **(Fig S6)**. Gene-set enrichment analyses were conducted on the MAGMA and STRING platforms, which identify 36 gene sets significantly enriched on at least one platform (FDR<0.05) **(Table S4)**. Most hsCRP-associated genes are involved in immune regulation and/or cholesterol metabolism. Based on the ClueGO pathway network analysis, 55 pathways were statistically significant (*P_Bonferroni_*<0.05) and were clustered into five functional groups **(Fig 3)**. The representative pathways of five groups suggested that hsCRP levels may be involved in cholesterol metabolism (“cholesterol metabolic process” and “acylglycerol homeostasis”), immune process (“cellular response to virus” and “CD4-positive, alpha-beta T cell differentiation involved in immune response”), and endocrine system (“endocrine pancreas development”). Among all expression quantitative trait loci (eQTL) for hsCRP lead SNPs and gene expression in liver and blood tissues **(Table S5)**, a total of 13 pairs of SNP-gene remained statistically significant in the liver (*P*<2.25×10^-8^) and blood (*P*<2.46×10^-8^) tissues after the Bonferroni correction. Notably, some eQTLs were linked to distant genes beyond their nearest genes (i.e., rs4486443 in *ADAR* and rs1260326 in *GCKR*).

**Fig 2.**
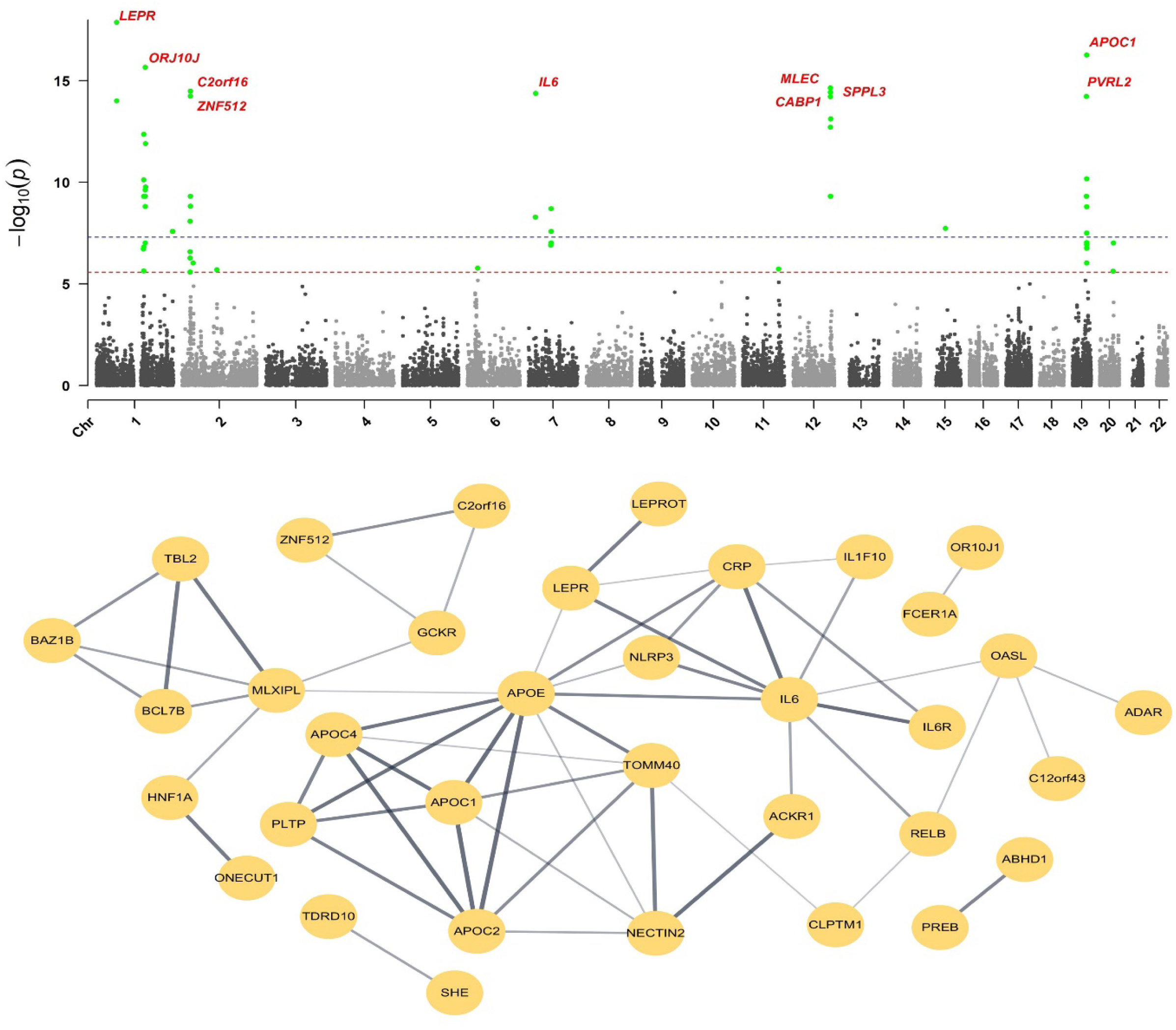
Gene-based association analysis of hsCRP. **(A) Manhattan plot of genome-wide gene-based analysis of hsCRP.** Red and blue dotted lines indicate the Bonferroni (*P*<2.68×10^-6^) and nominal genome-wide significance level (*P*<5×10^-8^). Labelled genes are the top 10 independent genes significantly associated with hsCRP. **(B) Interaction network between the genes associated with hsCRP.**

**Fig 3.**
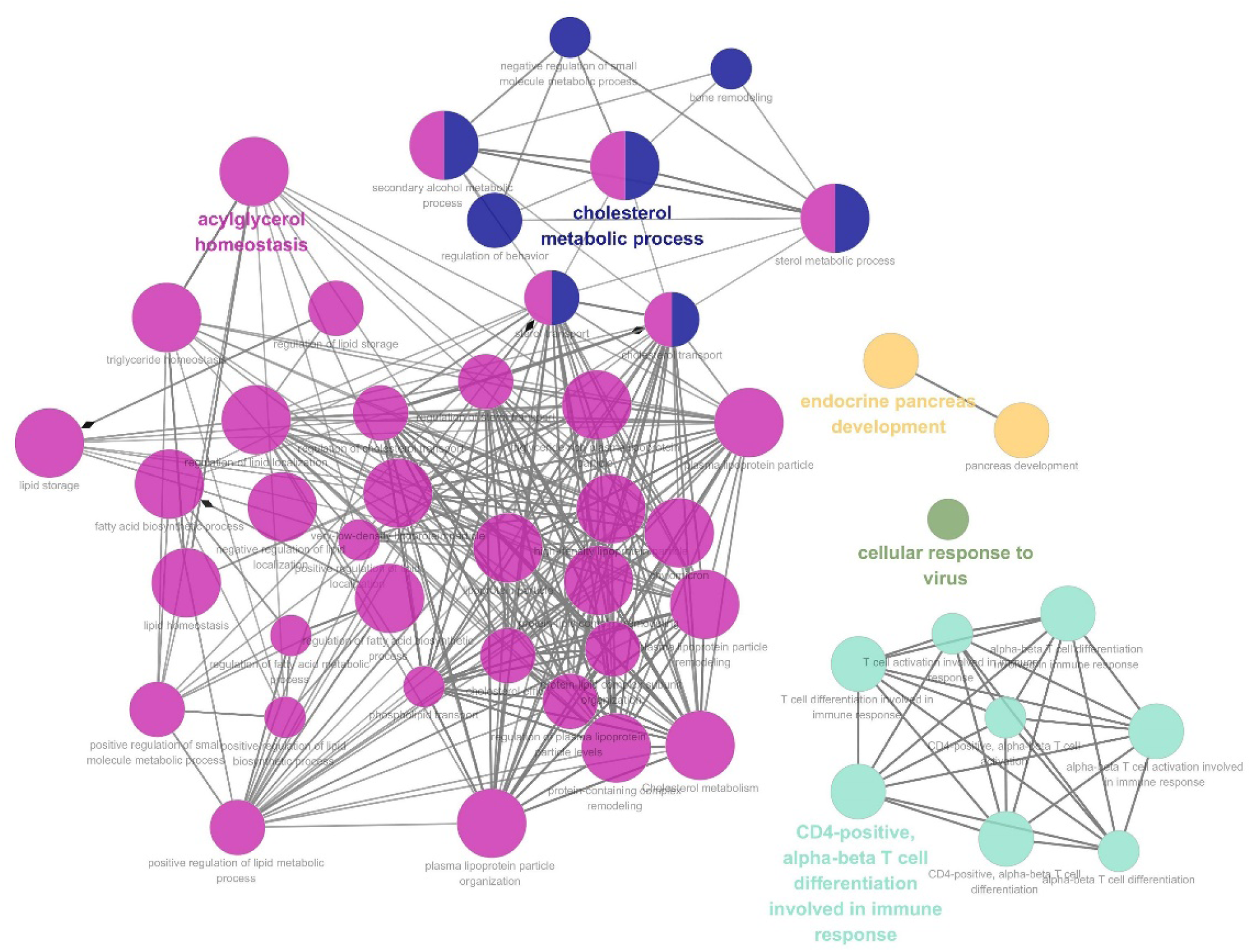
Association network of the hsCRP-associated pathways. This network depicts the association between pathways from gene ontology (GO) and the Kyoto Encyclopedia of Genes and Genomes (KEGG). The sizes of the nodes reflect the enrichment significance of the pathways. Color-labeled pathways represent the functional groups with the most significant *P*-value.

Furthermore, *PVRL2* showed the greatest number of eQTL pairs with the lead SNPs.

### Genetic relationships between hsCRP and other phenotypes

To investigate potential relationships between hsCRP levels and other diseases/traits, PheWAS was conducted using hsCRP-polygenic risk score (PRS). **Fig S7** illustrates the overall distribution of hsCRP-PRS calculated from 71,019 participants, which approximates a standard normal distribution. A result of multivariable linear regression on hsCRP levels indicated that as a single-unit increase of hsCRP-PRS, the log-transformed hsCRP levels increased as 0.21 mg/L **(Table S6)**. A total of 65 continuous variables and 49 binary variables were evaluated for their relationships with hsCRP-PRS **(Table S7-8)**. As a result, a total of nine continuous variables were significant after the Bonferroni correction (*P*<7.69×10^-4^=0.05/65 traits): LDL (β=−1.47, *P*=1.69×10^-33^), total cholesterol (β= −1.15, *P*=1.59×10^-17^), platelet count (β= 1.34, *P*=7.64×10^-8^), γ -glutamyl transpeptidase (γ-GTP, β=0.84, *P*=8.30×10^-8^), uric acid (β=0.02, *P*=2.41×10^-7^), height (β=−0.08, *P*=5.18×10^-5^), weight (β=−0.06, *P*=1.21×10^-4^), fibrinogen (β=1.99, *P*=2.19×10^-4^), and mean corpuscular volume (β=−0.07, *P*=5.81×10^-4^) **(Fig S8, Table S7)**.

LDL level is a classic modifiable risk factor for CVD along with hsCRP (4), therefore, we investigated the genetic association between LDL and hsCRP levels. We found a positive association between the actual levels of LDL and hsCRP. Specifically, as LDL cholesterol levels increased, hsCRP levels also increased, so did the proportion of subjects with elevated hsCRP levels (≥3 mg/L) **(Fig 5a)**. In contrast, hsCRP-PRS and LDL levels showed an inverse trend **(Fig 5b)**. As the LDL cholesterol levels increased, the hsCRP-PRS levels decreased remarkably.

**Fig 4.**
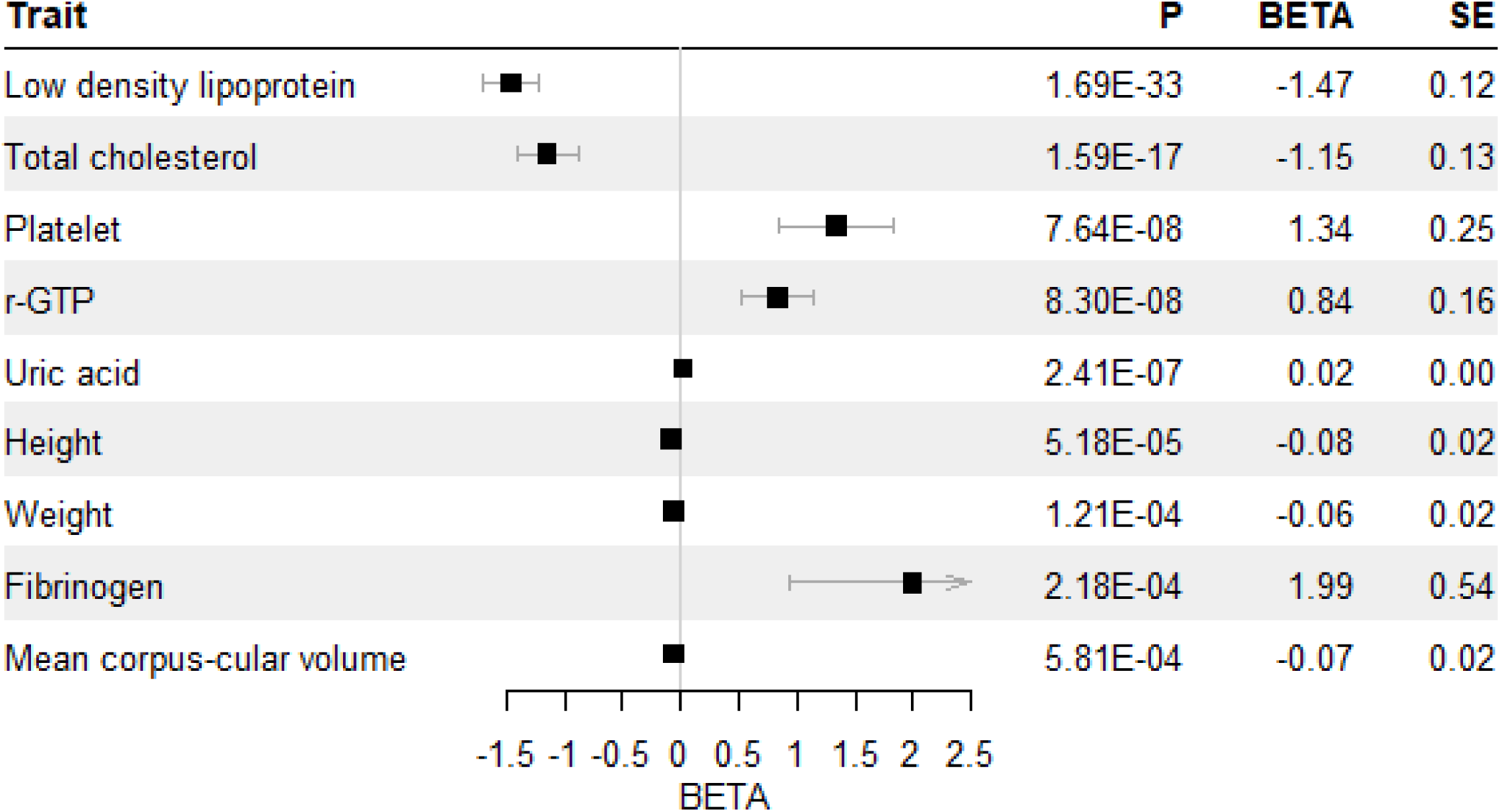
Forest plot of PheWAS using hsCRP-PRS. Bonferroni significance level of PheWAS *p*-value is 4.35×10^-4^.

**Fig 5.**
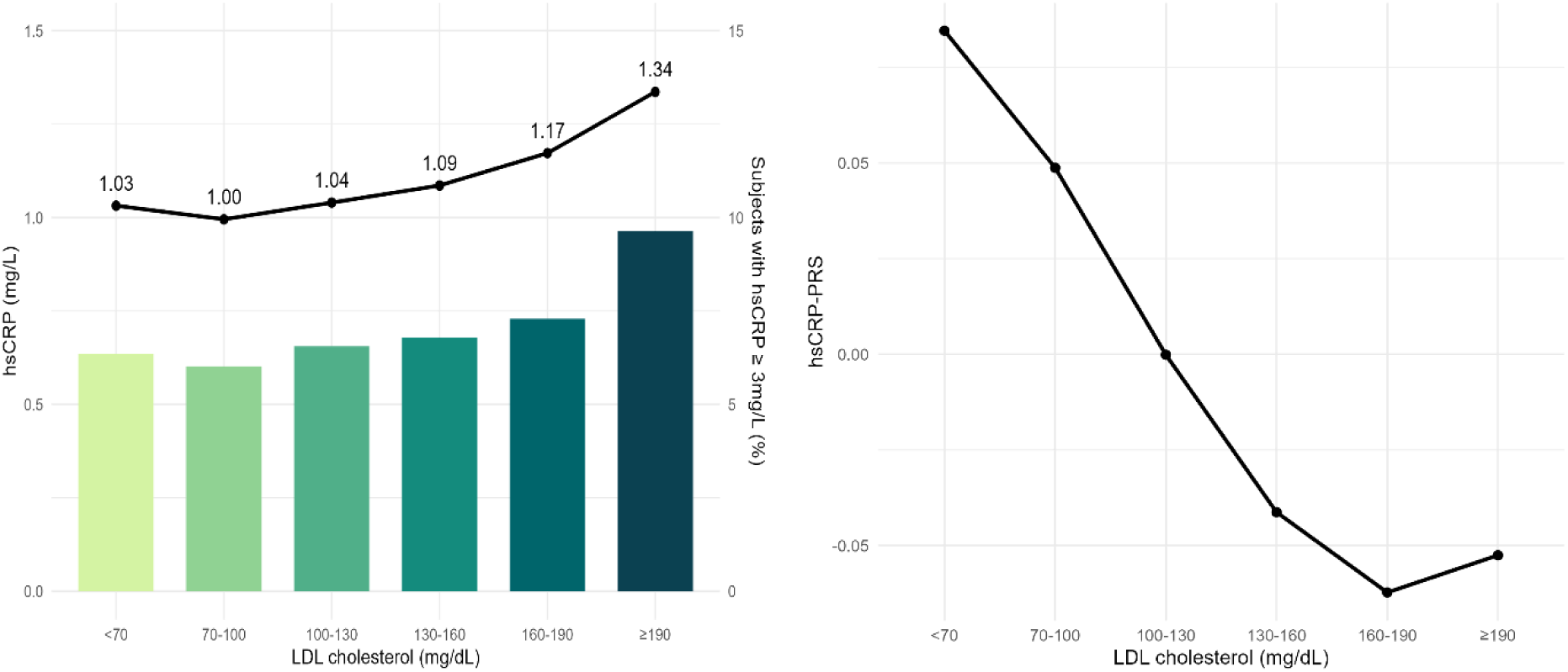
The association analyses between hsCRP and LDL. **(A) The association between hsCRP and LDL cholesterol.** A line graph indicates the hsCRP levels, and a bar graph indicates the proportion of subjects with elevated hsCRP levels. **(B) The association between hsCRP-PRS and LDL cholesterol.**

## Discussion

Inflammation is associated with many chronic diseases (1–4). As a key biomarker of systemic inflammation, hsCRP has been suggested as a risk factor for multiple diseases (5, 6). CVD is strongly associated with hsCRP levels (1, 4). Although LDL is deemed as a causal factor of CVD, recent analyses reported that high hsCRP levels are associated with CVD risk despite low LDL levels (27). Considering the clinical importance of hsCRP, we performed a GWAS on hsCRP in 71,019 Koreans and identified 69 lead SNPs, including 12 novel variants. In the current study, 10.15% of the variance in hsCRP levels could be explained by 69 lead SNPs. Furthermore, 58 hsCRP-associated genes have been identified using genome-wide gene-based analyses. These genes are primarily involved in the immune system and cholesterol metabolism. Subsequent PRS-PheWAS analysis showed a concordant trend, indicating that genetically determined hsCRP was associated with LDL, total cholesterol, and platelet counts. Specifically, we highlighted its association with LDL, which is a risk factor for CVD along with hsCRP (4).

As this is the largest hsCRP-GWAS conducted in a Korean population, the identified SNPs may provide insights into the genetic factors associated with chronic inflammation in EA populations. The most significant SNP was rs3093068 in *CRP* (β(C)_meta_=0.246, *P*_meta_=1.28×10^-304^), and *PVRL2* was the gene harboring the largest number of lead SNPs (4 SNPs). Although *CRPP1* is a pseudogene of *CRP*, evidence indicates that it may function as an enhancer of *CRP* by regulating prolonged transcription (28). Further validation analysis conducted on SNPs across the three groups (EA ancestry, multi-ancestry with EA, and multi-ancestry without EA) demonstrated that the SNPs identified in the EA ethnic populations were validated better than the others (EA ancestry: 51.5%, multi-ancestry with EA: 29.2%, and multi-ancestry without EA: 36.8%). This suggests that differences in the genetic architecture between populations of different ancestry may influence susceptibility to hsCRP SNPs. This evidence strengthens the importance of hsCRP genetic analyses in diverse populations.

The most significant 3 hsCRP-associated genes were *LEPR* (*P*=1.38×10^-18^), *APOC1* (*P*=5.55×10^-17^), and *OR10J1* (*P*=2.22×10^-16^). *LEPR* encodes a leptin receptor protein. Leptin is an adipocyte-derived hormone that regulates food intake and energy metabolism. Leptin stimulates hepatic and vascular CRP synthesis (29). In the current study, there were two lead SNPs in *LEPR*; rs75276031 (β(T)_meta_=0.13, *P*_meta_=4.54×10^-11^) and rs6588147 (β(A)_meta_=0.06, *P*_meta_=9.81×10^-17^). According to a previous protein quantitative trait study, the minor allele of rs6588147 is associated with increased levels of leptin receptor protein (30). This may promote the actions of leptin, including CRP synthesis. In addition, *APOC1* encodes the ApoC1 protein, which is known to modulate the metabolism of high-and very low-density lipoprotein (31). In an immunity perspective, ApoC1 was reported to strongly bind to lipopolysaccharide (LPS) of gram-negative bacteria, stimulating the LPS-induced TNFα production by macrophages (32). Systemic inflammation may also be linked to elevated hsCRP levels. *OR10J1* encodes olfactory receptors. Although it might seem irrelevant, various types of olfactory receptors are expressed in macrophages of both mice and humans. Although their exact functions are unknown, olfactory receptors in macrophages participate in endogenous oxidation and metabolic processes that may contribute to inflammation control (33). Furthermore, several GWAS studies suggested that some SNPs in *OR10J1* were associated with CRP levels (34, 35). In addition to the top three implicated genes, most hsCRP-associated genes were mainly involved in cholesterol metabolism (*APOC1*, *APOE*, *APOC4,* etc.) and the immune response (*IL6*, *IL6R*, and *IL1F10*). This was strengthened by subsequent analyses (gene set and pathway network analyses), which highlighted the link between hsCRP levels and cholesterol or immunity pathways. This indicates that hsCRP levels may be elevated in the presence of variants or mutations in genes involved in cholesterol metabolism or immunity. This is consistent with previous findings that serum CRP regulates the immune response by interacting with cholesterol in the blood vessels (4). However, in the current study, cholesterol-related genes or pathways tended to be more enriched than in other GWAS on hsCRP (11, 14, 36).

To evaluate the effects of genetically determined hsCRP levels on other traits, a PRS-PheWAS analysis was conducted. There were nine clinical traits associated with hsCRP-PRS, of which five (γ-GTP, LDL, total cholesterol, platelet, and fibrinogen) were remarkable. The serum γ-GTP has been highlighted as a marker of oxidative stress independent of the obesity (37). Considering that oxidative stress and inflammatory response promote each other(38), elevated serum γ-GTP may be associated with the inflammation caused by oxidative stress (37). Several *in vitro* studies have indicated that chronic inflammation could decrease LDL levels by promoting *LDLR* expression (39) but could also increase susceptibility to oxidation (40). It is more atherogenic, amplifying the inflammatory response and accumulation of cholesterol in macrophage (41). Platelets are known to play a role in inflammatory response and immune regulation beyond hemostasis (42). During chronic inflammation, platelets interact with white blood and endothelial cells and release cytokines and chemokines that promote inflammation (42, 43). Fibrinogen is a glycoprotein synthesized by hepatocytes and plays a key role in the formation of thrombus, the most common cause of CVD (44). Moreover, a growing body of evidence indicates that fibrinogen is an indicator of proinflammatory state (45). Based on the overall PRS-PheWAS results, an increase in genetically determined hsCRP levels may contribute to a decrease in cholesterol levels and a stronger oxidative environment in the blood vessels, which can further promote inflammation. Thus, individuals with higher hsCRP-PRS levels may be more susceptible to CVD owing to their oxidative susceptibility, even when LDL levels are low. This suggestion can be partially reinforced by the previous reports that those with low LDL (<130 mg/dL and <70 mg/dL) and high hsCRP levels (≥2 mg/L) had higher CVD risks than those with high LDL and low hsCRP levels (46, 47).

This study has some limitations. First, because GWAS identify variants associated with complex phenotypes (14), hsCRP susceptibility SNPs may not be the causal variants of hsCRP levels. Therefore, we focused on uncovering the underlying biological pathways and functions, rather than on functional implications of individual SNPs. Second, chronic inflammation is a complex condition that may be influenced by various factors. Therefore, residual confounding factors may not have been controlled for. Third, the highlighted genes and pathways were identified by *in silico* analysis. Further *in vitro* and *in vivo* functional investigations are required to confirm the underlying biological mechanism of hsCRP. Fourth, since PheWAS was conducted in the same population as GWAS, there might be a risk of overfitting when investigating the association between hsCRP-PRS and other traits.

This study is one of the largest GWAS on hsCRP levels in the EA population. A total of 69 independent hsCRP susceptibility SNPs were identified, of which 12 were novel. Subsequent analyses highlighted the relationship between hsCRP SNPs and genes/pathways associated with cholesterol metabolism and immune responses. Notably, alleles associated with higher hsCRP levels in the form of PRS were also associated with lower cholesterol levels, in contrast to observational epidemiological studies. The findings of the present study reinforce the clinical importance of hsCRP levels and hsCRP-PRS may be a clinical tool for predicting the potential risk of abnormal cholesterol metabolism and its related diseases.

## Materials and Methods

### Study population and data collection

This study was conducted using bioresources from the National Biobank of Korea and Center for Disease Control and Prevention, Republic of Korea (NBK-2022-004). In this study, indpendent subjects were recruited from three population-based KoGES cohorts (22); HEXA, CAVAS, and the Ansan/Ansung study. Among the 58,694 HEXA, 8,105 CAVAS, and 8,840 Ansan/Ansung participants, those without genetic data (N_HEXA_=1; N_Ansan/Ansung study_=3,347) were excluded. Those without hsCRP data were excluded (N_HEXA_=5, N_CAVAS_=860, N_Ansan/Ansung study_=15), as well as those with hsCRP level above 10 mg/L (N_HEXA_=254, N_CAVAS_=137, N_Ansan/Ansung study_=1) because high CRP levels could be caused by acute inflammation (48). In the case of an outlier or missing value for hsCRP level, hsCRP data from the closest follow-up study were used if available (i.e., the 1^st^ follow-up study for HEXA and the 1^st^ to 8^th^ follow-up study for the Ansan/Ansung study) to minimize exclusions. Finally, 71,019 study participants (N_HEXA_=58,434; N_CAVAS_=7,108; N_Ansan/Ansung study_=5,477) were selected for the analysis **(Fig S1)**.

Demographic and epidemiological data were collected using self-reported questionnaires, and laboratory experiments for clinical data were conducted in hospitals across Korea. CRP levels were measured using a high-sensitivity hsCRP method. We collected data on age, BMI, sex, alcohol consumption, smoking, regular physical activity, family income, educational attainment, and chronic disease history simultaneously with hsCRP level collection. Phenotypic data for PheWAS were extracted from KoGES HEXA, CAVAS, and Ansan Ansung databases. Among the available epidemiological and clinical variables, those with valid values exceeding 10,000 or with inflammation-related phenotypes were selected. Among these, binary variables for which the observed frequency was less than 20 in each category were excluded. In total, 114 phenotypes (e.g., platelets, LDL, and total cholesterol) were detected for PheWAS **(Table S7-8)**. The study protocol was approved by the Korea National Institutes of Health and the Institutional Review Board (IRB: CBNU-202306-HR-0153) of Chungbuk National University.

### Genotyping and quality control

Using the Korea Biobank Array (KCHIP) platform, approximately 800,000 (800 K) SNPs were genotyped from each sample (49). Initially, quality control (QC) procedures were performed to exclude individuals from further analyses whose samples met at least one of the following criteria: 1) call rate<97%, 2) excessive singletons, 3) sex discrepancy, 4) cryptic second-degree relatives, or 5) withdrawal and blind replicates [20]. SNP QC was also conducted, excluding SNPs with 1) Hardy-Weinberg Equilibrium (HWE) *P<*1×10^-6^, 2) call rate<0.95, or 3) low quality SNP (variants classified into another category by R package SNPolisher or off-target variants), resulting in approximately 460 K SNPs remaining (49). Imputation was performed using IMPUTE 4 software based on the Korean reference genome (49). Briefly, 8,000K SNPs were tesed after the following exclusion criteria were met: 1) information content metric<0.8, and 2) minor allele frequency (MAF)<0.01 (49). For the current GWAS, we double-checked and excluded genotyped or imputed SNPs or samples according to the following criteria: 1) MAF<0.01, 2) HWE *P*<1 × 10^-6^, 3) missingness per individual<0.05, and 4) missing data per marker<0.05. Consequently, 7,981,597 genotyped or imputed SNPs for HEXA; 7,971,416 for CAVAS; and 7,972,085 for the Ansan/Ansung study were used in the current analysis.

### Statistical analysis of epidemiological variables

The hsCRP levels were considered continuous; however, owing to the positively skewed distribution, the natural log-transformed hsCRP levels were used for the current analyses. Among the demographic and epidemiological variables, age and BMI were analyzed as continuous variables; sex, alcohol drinking, smoking, regular physical activity, and history of chronic disease were analyzed as binary variables; and family income and educational attainment were categorized into three or four groups, respectively. To investigate the associations between hsCRP levels and potential confounding variables, several statistical analyses were conducted according to the different types of variables. Student’s t-test and one-way analysis of variance were used to compare hsCRP levels between two groups and among three or more groups of categorical variables, respectively. Pearson’s correlation test was used for continuous variables. We initially constructed a full model using the multivariable linear regression for hsCRP (as dependent variable) with all potential confounding variables statistically significant in the three KoGES cohorts in common (*P*<0.05, as independent variables), and then backward selection was performed to extract the covariates. Accordingly, age, sex, BMI, regular physical activity, smoking, and history of hypertension were controlled for in subsequent analyses. In addition, we examined the relationship between LDL and hsCRP levels, both of which are commonly used as biomarkers of CVD. Participants were categorized based on the Korean clinical guidelines for LDL levels:<70 mg/dL, 70–99 mg/dL, 100–129 mg/dL, 130–159 mg/dL, 160–189 mg/dL, and >190 mg/dL (50). Furthermore, hsCRP levels, proportions of subjects with elevated hsCRP (≥3 mg/L), and hsCRP-PRS were compared according to the LDL level categories. All statistical analyses were performed using R Software (version 4.3).

### GWAS on hsCRP levels

For the GWAS of hsCRP levels, a multivariable linear model with adjustments, as explained above, was used. A GWAS of hsCRP levels was performed in each of the three KoGES cohorts (HEXA, CAVAS, and Ansan/Ansung study) using PLINK (version1.9) (51). To combine the results of all three KoGES cohort studies, we conducted a fixed-effects inverse variance-weighted meta-analysis using METAL (52), as the overall variation due to heterogeneity was relatively low (I^2^=14.65%) (53). For a genomic control, the statistics of each GWAS result were corrected by the METAL based on the genomic inflation factors (λ_HEXA_=1.09, λ_CAVAS_=1.01, and, λ_Ansan/Ansung study_=1.01). With these corrections, the inflation factor for the meta-analysis was 1.02, indicating little systemic inflation (54).

Human leukocyte antigen (HLA) variants are relatively few, but their complicated LD structure could make GWAS results difficult to interpret (55), we excluded genetic variants in the HLA regions (chr6:25Mb-33Mb, hg19) from further analyses. Furthermore, SNPs on the sex chromosome and indels were excluded because they differed according to sex or individual. We considered hsCRP susceptibility SNPs that satisfied the nominal genome-wide significance threshold (*P*<5×10^-8^) in at least one of the three KoGES cohorts, with successful validation (*P*<0.05) in the remaining KoGES cohorts, and with nominal genome-wide significance in the combined results generated by the meta-analysis (*P_meta_*<5×10^-8^). Using the extracted hsCRP susceptibility SNPs, we performed LD clumping using FUMA platform (23). The GWAS-based hsCRP susceptibility SNPs were grouped into the independent loci based on the distance (±500kb) and LD structure (r^2^<0.1). For each independent genetic locus, the SNP with the smallest *P_meta_* value was indexed as the lead SNP.

To calculate the proportion of hsCRP variance explained by the lead SNPs, the following formula was used (14);

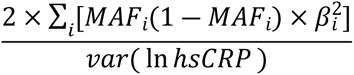

In this case, ∑ is the sum, *MAF*_*i*_ is the MAF of the corresponding lead SNP*i*, *β*_*i*_ is the effect estimate of the SNP*i* on natural log hsCRP level, and *var* is the variance of natural log hsCRP levels.

### Validation of hsCRP susceptibility SNPs identified by previously published GWAS

Previous hsCRP GWAS summaries were obtained from the GWAS catalog on 08/31/2023 (56). To consider possible ethnic differences in genetic susceptibility loci, EA and with/without EA validation analyses were conducted (number of evaluated SNPs: N_ancestry (EA)_: 253, N_multi-ancestry with EA_: 2,485, N_multi-ancestry without EA_: 2,232). SNPs on the sex chromosome or with *P*>5×10^-8^, as well as indels, were excluded from validation. Variants that were not used in our GWAS were also excluded. For hsCRP susceptibility SNPs that have been reported several times, those with the smallest *P*-value were selected. The remaining SNPs were clustered based on distance and LD structure (ancestry: ±500kb; r^2^<0.5, multi-ancestry (with/without EA): ±10,000kb; r^2^<0.001). As a result, 123, 260, and 239 SNPs were evaluated in the ancestry and multi-ancestry (with/without EA) validation analyses, respectively. SNPs were considered significantly validated when *P*_meta_<0.05, and the results of the previous and current studies were concordant. The detailed process is shown in **Fig S2**.

### Functional analysis

Based on the significant hsCRP SNPs from the meta-GWAS results (*P*_meta_<5 × 10^-8^), we performed the ANNOVAR enrichment test using FUMA software (23), where SNPs were functionally annotated as either exonic, intronic, intergenic, splicing site, 5′/3′-UTR, or upstream/downstream of genes, presented in **Fig S5** (57). Subsequently, functionally annotated SNPs were mapped to genes using two strategies: positional mapping and eQTL mapping (23). Positional mapping was based on the physical distance between the SNPs and genes within a maximum distance of 10Kb. The eQTL mapping was conducted based on *cis*-eQTL association between SNPs and genes, limiting the distance to 1Mb. eQTL data were obtained from Genotype-Tissue Expression (GTEx) version 8 (58), and the Bonferroni-corrected significance level was used based on the number of genes tested for each tissue.

Using MAGMA (24), we performed gene-based, gene-set, and gene-property analyses based on the overall summary results of the meta-GWAS on hsCRP. Through gene-based analyses, individual SNP statistics within protein-coding genes were combined, resulting in gene-based statistics. The genome-wide significance level of the gene-based analysis was considered using the Bonferroni correction (*P*<2.68 × 10^-6^=0.05/18,681 genes). Using gene-set analysis, we explored which genes in a specific gene set were more strongly associated with hsCRP levels than other genes. The curated gene sets and gene ontology data obtained from the molecular signature database (MsigDB version 7.0) (59) were used as input data. Gene-based *p*-values from the gene-based analysis were used to compute gene-set statistics. A total of 15,480 gene sets were tested, and the significance level was set according to the false discovery rate (FDR) control (FDR<0.05). Gene-property analysis was performed to identify the tissue-specific expression of hsCRP-associated genes, and RNA-sequence data for 30 general tissue types were obtained from GTEx version 8 (58).

STRING was used to predict protein-protein interactions within the input genes or proteins (25). In this study, STRING analyses were performed based on MAGMA gene-based analysis results. We investigated how prioritized genes (*P*<2.68 × 10^-6^, N=58) interact with each other to influence hsCRP levels. We set the minimum confidence score at 0.4 (medium confidence) corresponding to the estimated likelihood that the association between proteins was true. The STRING gene set enrichment test was performed on all genes (N=18,681) used in the MAGMA gene-based analysis. Various resources were used to collect gene-set input data, including gene ontology (60), Reactome (61), disease ontology (62), COMPARTMENTS (63), Monarch (64), Uniprot (65), Pfam (66), Interpro (67), KEGG (68), TISSUES (69), and simple modular architecture research tool (70).

The ClueGO plug-in in Cytoscape was used to facilitate the biological interpretation of hsCRP-associated genes (26). The functional enrichment analysis was performed using MAGMA gene-based analysis results, as well as gene ontology and KEGG databases (60, 68). Pathways containing four or more hsCRP-associated genes were selected and their significances were calculated using a two-sided hypergeometric test with Bonferroni correction (*P_Bonferroni_*<0.05). Based on the kappa statics, the degree of linkage between pathways in network was estimated and functional groups were generated (kappa score≥0.4).

### PRS and PheWAS

Using the hsCRP-PRS constructed using lead SNPs, PheWAS was conducted to investigate the relationship between cumulative genetic factors of hsCRP and a variety of phenotypes. To compute the hsCRP-PRS, 69 lead SNP statistics were extracted, and the following formula was used:

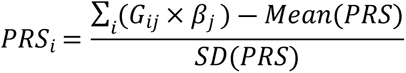

where *i* and *j* represent the individual subjects and SNP, respectively. *G*_*ij*_ is the number of hsCRP-increasing alleles of SNP*j*, and *β*_*j*_ means the effect estimate of SNP*j*. The *Mean*(*PRS*) and *SD*(*PRS*) refer to the mean and standard deviation of the PRS calculated from the 71,019 participants. PLINK (version1.9) was used for hsCRP-PRS calculation (51). PheWAS was conducted using the same covariates used in the hsCRP GWAS (age, sex, BMI, regular physical activity, smoking, and history of hypertension). Multivariable logistic regression models were adapted for binary variables (intestinal polyps, breast cancer, and family history of cancer), while multivariable linear regression models were adapted for continuous variables (height, LDL, and total cholesterol). PheWAS analysis was performed using the R package PheWAS (71).

## Acknowledgements

The authors thank all individuals who participated in this study.

## Supporting information captions

### Supporting information 1

**Fig S1.** Flow chart of study population selection

**Fig S2.** Flow chart of SNP selection for validation analysis

**Fig S3.** Manhattan plots of GWAS on hsCRP for three KoGES cohorts (A) Manhattan plot of GWAS on hsCRP for HEXA cohort (B) Manhattan plot of GWAS on hsCRP for CAVAS cohort (C) Manhattan plot of GWAS on hsCRP for Ansan/Ansung study cohort

**Fig S4.** Manhattan plot of meta-GWAS on hsCRP for three KoGES cohorts

**Fig S5.** Proportion plot of functionally annotated SNPs

**Fig S6.** MAGMA gene-property analysis result

**Fig S7.** Density plot of hsCRP-PRS

**Fig S8.** PheWAS plot of 114 traits from hsCRP weighted PRS

**Table S1.** Characteristics of the study participants

**Table S2.** Mean of hsCRP levels and associations with general characteristics

**Table S3.** Significant genes of MAGMA gene-based analysis on hsCRP GWAS

**Table S4.** FDR significant gene-sets according to the enrichment test in MAGMA and STRING

**Table S5.** Significant eQTL pairs between hsCRP lead SNPs and genes in the liver and blood tissues

**Table S6.** Multivariable linear regression model on log-transformed hsCRP

**Table S7.** PheWAS results for continuous variables

**Table S8.** PheWAS results for binary variables

### Supporting information 2

**Data S1.** SNPs associated with hsCRP by GWAS

**Data S2.** Validation of SNPs discovered in East Asian ancestry subjects

**Data S3.** Validation of SNPs discovered in multi-ancestry subjects containing East Asian

**Data S4.** Validation of SNPs discovered in multi-ancestry subjects not containing East Asian

